# Tumorcode - A framework to simulate vascularized tumors

**DOI:** 10.1101/216903

**Authors:** Thierry Fredrich, Michael Welter, Heiko Rieger

**Affiliations:** Theoretical physics, Department for natural sciences and technology, Saarland University, Germany

## Abstract

During the past years our group published several articles using computer simulations to address the complex interaction of tumors and the vasculature as underlying transport network. Advances in imaging and lab techniques pushed in vitro research of tumor spheroids forward and animal models as well as clinical studies provided more insights to single processes taking part in tumor growth, however, an overall picture is still missing. Computer simulations are a none-invasive option to cumulate current knowledge and form a quasi in vivo system. In our software, several known models were assembled into a multi-scale approach which allows to study length scales relevant for clinical applications.

We release our code to the public domain, together with a detailed description of the implementation and several examples, with the hope of usage and futher development by the community. Justification for the included algorithms and the biological models was obtained in previous publications, here we summarize technical aspects following the workflow of a typical simulation procedure.

## 1 Introduction

Vascular networks play an important role in therapy and progression of cancer. In particular their formation and modifications are a special feature of most malignant tumors [1]. In the case of solid tumors, the growth of cancer cells is facilitated by the diffusion of nutrients within the vicinity of the cells. Growing beyond the average diffusion length of nutrients, tumors adopt new strategies to acquire nutrients. By transforming the present blood vessels, the tumor exploits them for his own purposes. The full complexity of this process is not yet understood.

Since the vasculature transports drugs to cancer cells, this transformation is certainly of interest for chemotherapy, and also radio therapy is known to be dependent on the oxygen level inside the tissue which is again dependent on the vessel morphology and vessel wall properties. The complex dependence of solute transport or solute distribution and spatial arrangement of the blood vessels is therefore of great importance in every cancer therapy.

Obtaining experimental or clinical data of sufficiently high resolution to reconstruct vascular networks is by itself a hard task [2]. To overcome the lack of data, in silico modeling of vasculature is done by various authors. In [3] an off-lattice, agent-based model is used to successfully construct vascular structures of around 1mm in size. Others also focus on the extra cellular matrix [4], but most studies focusing on structural aspects neglect blood transport inside the vessels which is essential for clinical applications. Well established models consider the blood flow but are restricted to two dimensions [5] and only a few are done in 3D [6]. Apart from blood vessel network construction, the growth of tumors is of key interest.

The implementation strategies for tumor growth fall into two categories. First: discrete, particle based approaches where each cancerous cell is modeled individually [7].

Naturally this takes considerable computational efforts providing greater details on the cell level. Second: models where the tumor is described by continuous equations, reaching dimensions relevant for diagnostics [8]. Nevertheless, it is mostly neglected that tumors in patients or animal models grow within well-vascularized environments which means the original or initial network is not adequately represented.

Another problem in the field is that the code is maintained and used by corresponding groups only, rather than sharing implementations which would simply save time. Repositories for open source code are well known in computer science and started to become useful in life science as well [9] [10].

To create blood vessel networks, we use a lattice-based attempt which discretizes the angular freedom and enables us to explore large scales. To the best of our knowledge there is no other software constructing synthetic arterio-venous blood vessel networks on macroscopic tissue volumes in 3 dimension matching topological, morphological and hydrodynamic properties observed in real samples. Our multi-scale approach describes tumor growth by partial differential equations and thus it is possible to simulate system sizes of the order of centimeters.

Subsequent, the interpretation of clinical studies, animal models and testing of hypotheses on biological processes is feasible. Our approach allows us to study topological and functional aspects at the same time which is relevant for therapeutics.

Since the understanding of malignant tumors is of great interest in the life science community and in-silico experiments are an extension of traditional biological techniques, we make our code, together with a detailed user manual, available to the public.

## 2 Design and implementation

Like modern implementations we reassemble two most distinct aspects. On the one hand we use *python* together with by the *scipy* packages for an easy and straight forward scripting of scientific problems. On the other hand, computationally expensive tasks are delegated to a core written in *C++* for performance reasons. Following up, a pythonic interface to this core has been created with the help of *Boost.Python.* We decided to use the *’hdf5’*- file format [11] to store data since it is highly scale-able and supported by both programming languages used.

### 2.1 Computational core

Standard *C++* coding is used where the comments follow the *doxygen* notation enabling the user to retrieve information about dependencies and inheritance within the object-oriented kernel. Instructions on how to create the documentation can be found in the subfolder *’doc’.*

Examples for demanding main applications are 1) calculating the hydro- and hemodynamics and 2) solving partial differential equations. Both problems reduce to solving systems of sparse linear equations which we do by the *Trilinos* package [12].

#### 2.1.1 Hydro- and hemodynamics

Before determining hydro- and hemodynamics of the network, the topology itself is created by the vesselgenerator (see folder *’src/common/vesselgen’*). The idea follows a 3 dimensional generalization of the algorithm presented in [13]. An example output can be found in Fig 2 and a complete description of the algorithm is provided in [2, section 2.1]. We summarize the basic steps as follows: First, arterial and venous root nodes are placed on a lattice and blood pressure or blood flow rate is assigned. The phenomenological formula, equation (3) in http://doi.org/10.1371/journal.pone.0161267.s001, is well suited for this task. Next, segments are appended at random until arterial and venous trees meet. Fixing the radii of the capillaries, Murray’s law allows us to determine the radii of all mother vessels up to the root node level. Once the radii are known, we assume mass conservation at each vertex to get a matrix equation analogues to a Kirchhoff circuit in electrodynamics. Solving these sparse matrix equation provides blood pressure values and blood flow rates for every vessel. Subsequently, the formula for laminar flow allows to calculate the shear force acting on the wall. In a last step, the total shear force is optimized in a Monte Carlo manner which requires to solve the sparse matrix equation frequently. Capillaries are deleted and inserted as long as the total shear force is minimized and the spatial distribution of capillaries is maximized. At the end of the construction step we take the Fâhræus-Lindqvist and phase separation effect into account.

**Fig 2.**
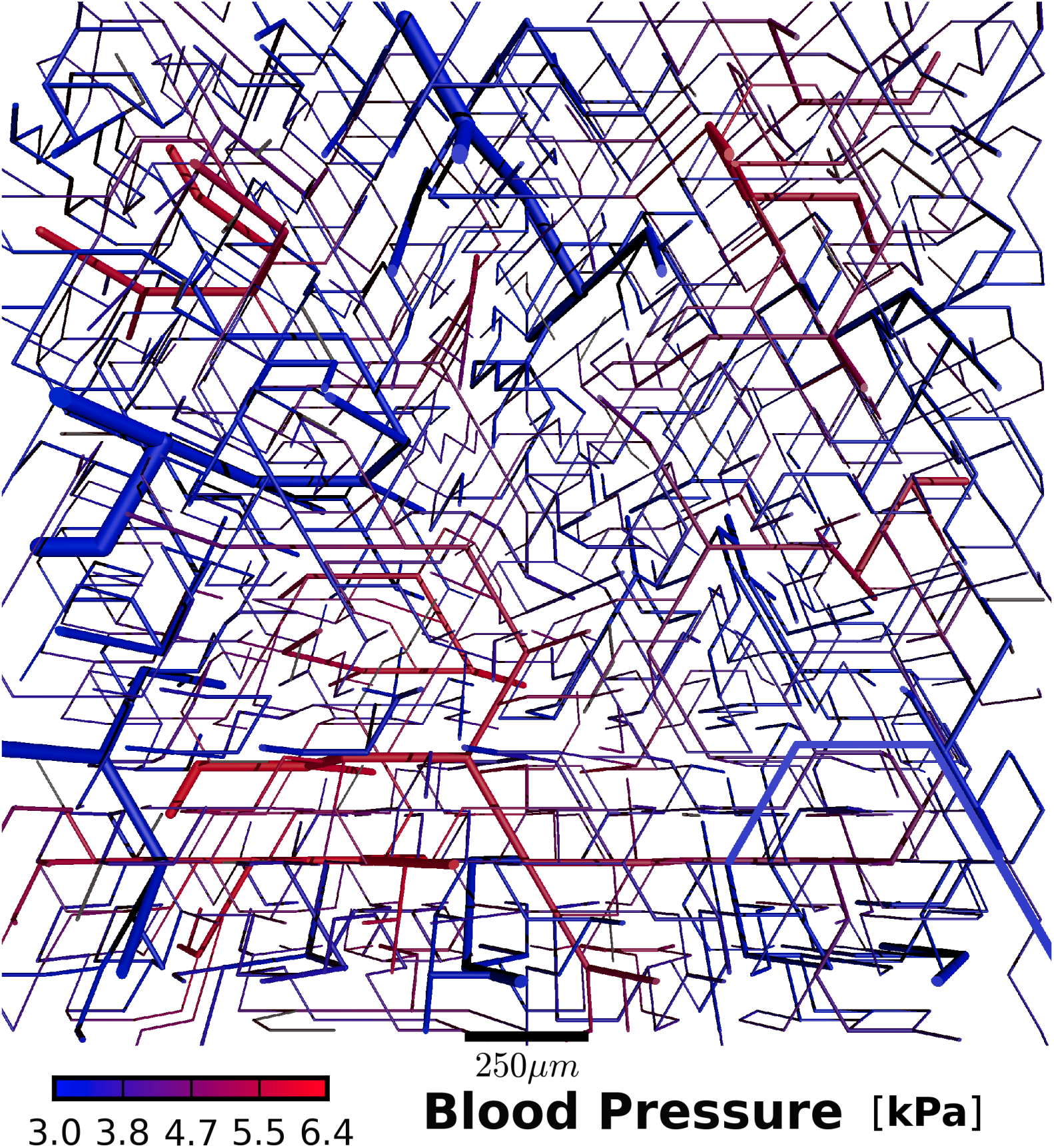
Rendering of blood vessel network. The rendering of the blood vessel network created in EX1 Blood vessel network creation is shown. Instructions on how to obtain such images from the software are explained in EX5 Visualization tools. Blood pressure is color-coded in *kPa* and a scale bar of 250 *μm* is present. A slice throughout the whole 3D cube is displayed in the figure.

#### 2.1.2 Tumor growth

The bulk tissue tumor model is based on a framework developed in [8], where the tissue is described as a mixture of various components. Our implementation includes normal, tumor and necrotic phases. The levelset-method is used to distinguish the different phases and defines the interface between them. Assuming incompressibility, one can describe the composition in terms of volume fractions which leads to a system of partial differential equations of diffusion-convection-reaction type for each phase. Introducing a regular cubic lattice for bulk tissue allows to solve these equations by Finite Difference techniques. Consequently, the numerical resolution of the *tumorcode* is governed by the lattice constant chosen. The number of grid points and vessels is limited by the provided hardware memory. Note that the bulk tissue and vascular network coexist in the same space, however, on different lattices. To fully utilize modern computing equipment we parallelized our code with the shared memory API of OpenMP. More details about this methodology could be found in the well written article [14].

For explorative studies, a fake tumor calculation which neglects the full dynamics of the cells and represents a tumor as spherically expanding mass of cells at a constant speed is implemented. Still the remodeling of the vasculature via vessel cooption, regression, and growth takes place.

#### 2.1.3 Finite Elements Method

In [15] we showed that the equation for the vascular oxygen distribution together with the diffusion equation for the tissue form a complicated nonlinear system of equations with the partial pressure values of oxygen as unknowns. For the longitudinal transport a bisection search is used to gather the partial oxygen pressure values within each vessel. Constructing consistent expressions for the transvascular exchange of oxygen across the vessel lumen was a challenging task, but enabled us to use the exchange as source or sink for the diffusion equation of the tissue oxygen field. Standard Galerkin methods were used to translate the task of solving the diffusion equation into solving sparse linear equations. To this end, the algebraic multigrid methods implemented in *Trilinos* were appropriate. Alternating the solver for vascular oxygen transport and the solver for the partial oxygen pressure of the tissue results in a stable self consistent equilibrium oxygen distribution.

### 2.2 Python API

After successful installation, the main tools can be found in the *’bin’* folder. Note that the Input Interface is written in *python* and that necessary parameters are also stored as python dictionaries (see folder *’py/krebsjobs/parameters’*).

*tumorcode* is ready to use on computing clusters equipped with a queuing system. To remember the fact that the computation will be submitted to a queuing system, the name of the simulation programs start with *submit*. Currently we support *’PBS (Portable Batch System)’* and *’Slurm Workload Manager’*. Job details such as expected runtime and allocated memory could be provided as optional commandline arguments. If no supported queuing system is present or the flag *(-q-local*) is invoked, the computation runs on the local machine. Possible commandline arguments and their structure are illustrated to the user by running the main programs with the help flag (*-h*). Each submit scripts expect the user to supply at least the name of a parameter set as command line argument.

How the single calculations are invoked and executed is presented in the supplemental material EX1 Blood vessel network creation to EX5 Visualization tools.

## 3 Workflow

The following subsections track a typical workflow corresponding to the yellow frames in Fig 1, where we visualize the procedure.

**Fig 1.**
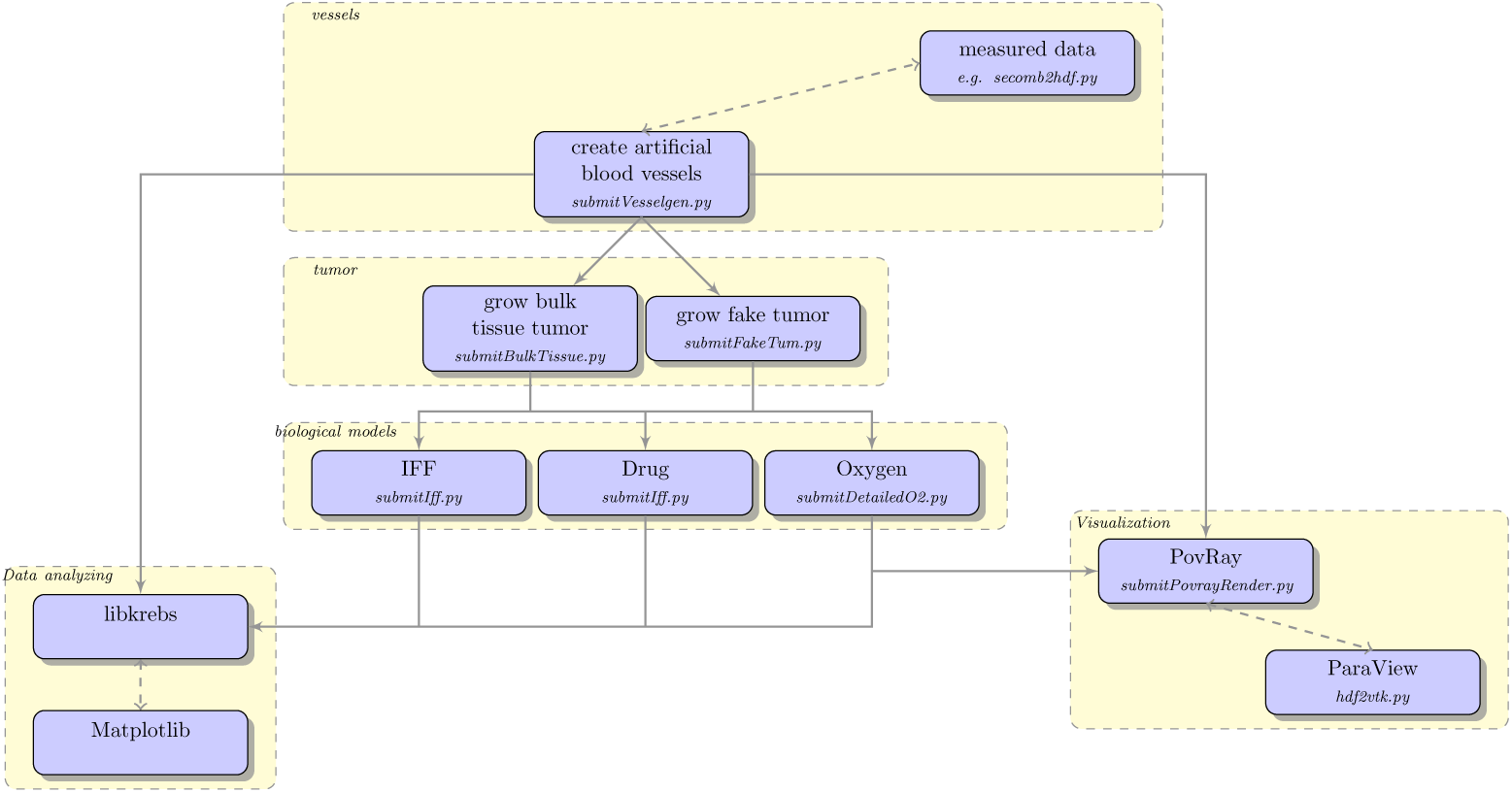
Workflow of *tumorcode*. Dotted lines indicate that connected modules can be interchanged. The straight arrows must be followed in the direction indicated by the arrow head.

### 3.1 Vessels

In general, one starts by creating synthetic blood vessels which serve as host and transport network (see 2.1.1). Another possibility is to provide measured input data. A transformation from the published data in [16] to the structure used by our software is described in the file *’py/krebs/adaption/apj2hdf.py’.* To elaborate the characteristic structure of a vessel file, a minimal working example is created where 3 arteries, 3 veins and 2 capillaries form a loop. In principal, each vessel is defined as an edge linking two nodes. Details can be obtained from Fig 4 as well as from the source stored in *’py/tests/simpleVesselConfig.py’*. The hemodynamic and morphological data calculated throughout the creation process as described in 2.1.1 is stored for each vessel within the ‘edges’ directory, e.g flow, radius and hematocrit. Further, the parameters used to calculate the hemodynamics, as specified by the *’-p’* flag, are written into the sub-folder ‘parameters’. The arrangement of root nodes accross the lattice is specified by the type ( *’-t’*) flag. The choice between 0 and 8 corresponds to one of the distribution shown in http://doi.org/10.1371/journal.pone.0161267.g003. The flag *(’-twoD’*) is of particular interest here. If selected, the creation process is restricted to a plane. All other programs are in full 3D.

**Fig 4.**
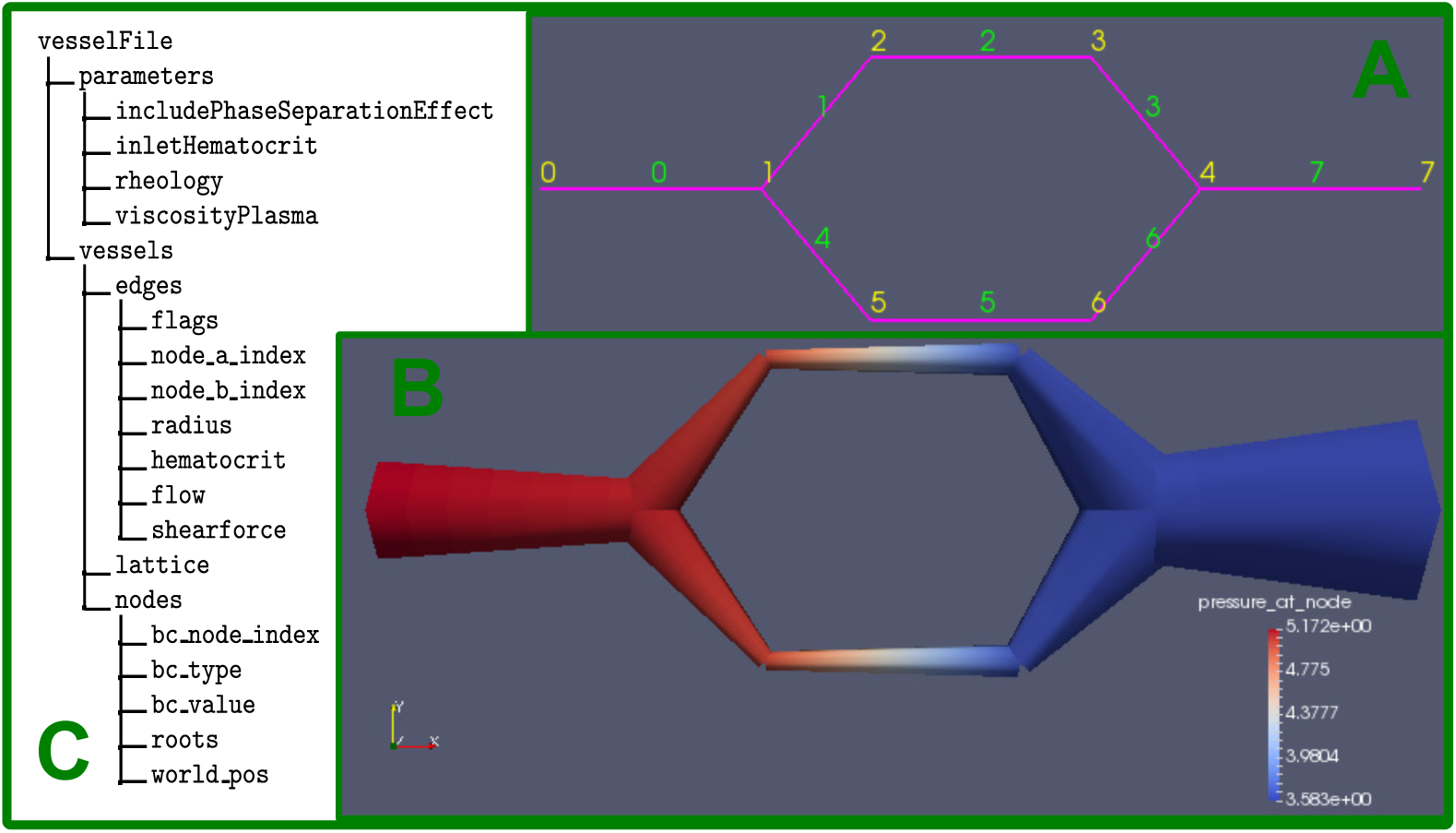
Illustration of vessel data structure. This minimal example contains 3 arteries, 3 veins and 2 capillaries. Inset A: the topological structure given by labels of the node points in yellow and the label of the edges in green. Inset B: morphology calculated by tumorcode, note the size radii of the vessels and the colorcode due to the pressure given in *kPa*. Inset C: structure of the *’hdf5’* file containing the vessel structure.

### 3.2 Tumor

Once vessels are constructed or read from experimental data, the workflow continues to the tumor stage where the simulation of either a fake tumor or a bulk tissue tumor can be started. *’submitBulkTissue’* runs simulations using the algorithm from the core discussed in 2.1.2 and *’submitFakeTum’* includes the discussed simplifications. Both type of tumor simulation need a parameter set and a hdf5 file containing proper vessels as command line input. When running, these simulations output their current configuration at time intervals as specified by the parameters. The groups in the output file are named out0000, out0001, etc. Fig 3 shows a snapshot of the simulation carried out in the supplemental material EX2 Tumor Growth.

**Fig 3.**
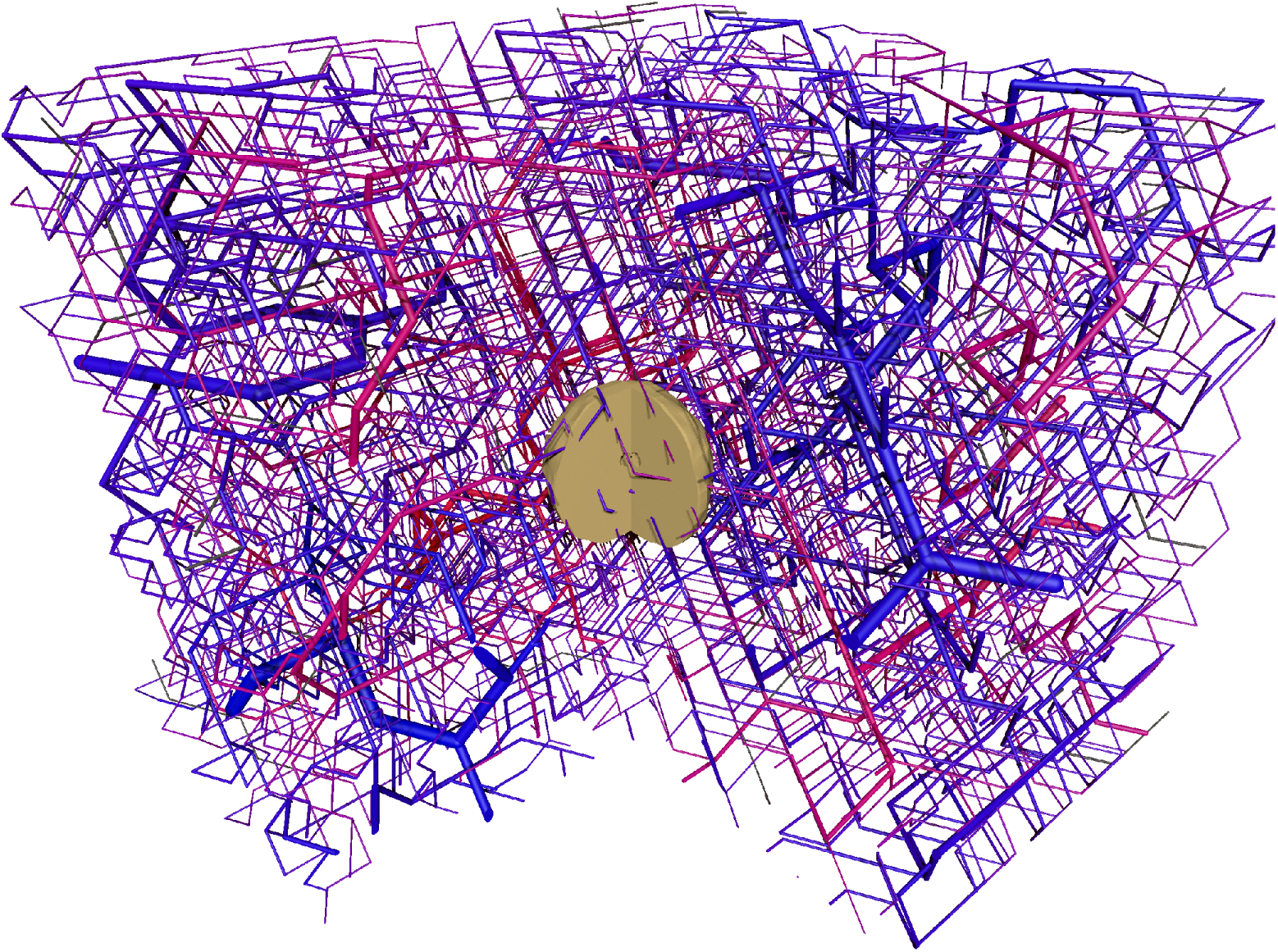
Rendering of a bulk tissue tumor simulation. Supplemental EX2 Tumor Growth explains how to obtain this image. The camera position ‘pie’ is chosen in order to see the tumor inside the cube. The vessel network is the same as shown in Fig 2

### 3.3 Biological Models

After the tumor stage, a *’hdf5*’-file containing time snapshots of either tumor simulation should be present. To invoke one of the biological models, the time point needs to be passed as additional argument. Therefore one needs to provide the name of a parameter set, the tumor file and the name of the time point. Note that time points are labeled consecutively by out0000, out0001 etc.

*’submitlff’* allows you to calculate distributions of the interstitial fluid flow and corresponding advection maps of drugs. Valid parameter sets can be found in *’/krebsjobs/parameters/parameterSetsIff.py’.* The sub-dictionary “iff” adjusts the interstitial fluid while the “ift” sub-dictionary adjusts the treatment of the tumor by the chemotherapy. Currently we provide 2 administration protocol.

**DF_INJECT_MODE_EXP** decreases the drug within the bloodstream in an exponential fashion. We correlate this with the injection of a single bolus decaying in the bloodstream.

**DF_INJECT_MODE_JUMP** let the level of drug within the blood stream jump from a constant value to zero. This mimics permanent infusion of a drug.

Varying the diffusion constant of the drug and its vessel wall permeability allows to screen for different modalities (see the supplemental EX3 Interstitial fluid flow and drug transport and the mentioned references).

Secondly, an elaborated model of blood and tissue oxygenation was developed [15]. Similarly to *’submitlff’, ‘submitDetailedO2’* needs a parameter set, a tumor simulation and a specified time point as input.

### 3.4 Analyzing and visualization

Finally, we offer a large number of scripts that we use to analyze raw simulation data.

Like the simulations, our analysis programs are hybrids between *C++* and *python.* Since the datasets can become quite large in filesize and take a long time to be processed, most evaluation programs are equipped with datacache i.e. the results of certain costly computations are saved to disk and loaded from there next time the execution is called with the same arguments. This facilitates iterative development, at least to some degree, without recomputation.

In order to use state of the art visualization techniques, we export datasets created with *tumorcode* in 2 different data formats as demonstrated in EX5 Visualization tools. *ParaView* allow an elaborated inspection of the vessel network structure. It offers interactive viewing and basic statistics on medium size systems, depending on your machines RAM. To render even the largest systems, we follow the ray trace approach of *POV-Ray.* Similar to *ParaView*, we offer a wrapper which converts vessel, tumor, oxygen and drug datasets to *POV-Ray* scene files, and run *POV-Ray* automatically in succession. Images of these kind are depicted in our latest publications [15,17].

## 4 Applications

As an illustration of the capabilities of *tumorcode*, we designed the supplemental examples EX1 Blood vessel network creation, EX2 Tumor Growth, EX3 Interstitial fluid flow and drug transport, EX4 Detailed oxygen distribution and EX5 Visualization tools within the realm of the two studies already published. However doing the same, we scaled the simulation in the examples to be smaller and quickly reproducible. On a standard desktop PC (Intel i7,2x8GB RAM) they are executed within minutes.

For details on the exact setup we refer to [15,17].

### 4.1 Blood vessels and tumor morphology

In silico modeling features the opportunity to measure structural and dynamical quantities at the same time. Simple averages of e.g. flow, radii and vessel volume are implemented straightforward in our software. Additionally, we can average data over spherical surfaces to mimic various experimental measurements. A profound inspection of a bulk tissue tumor simulation on a large scale could be found in [17, Fig 3]. There the graphs display microvessel density (MVD), vessel radius, cell velocity, oxygen, wall thickness and wall shear stress where the time course of tumor growth is visualized by different colors. The level set method allows to localize the distance of the tumor-tissue interface which is used as ordinate for the cited plots. Within that framework it is easy to define “inside” and “outside” of a tumor and focus on the differences.

Distinct experimental techniques such as MRI, micro-CT or xenografts are available to access morphological data of tumor tissue as well. But comparable data obtained from animal models usually requires the immolation of an animal per time point. Therefore such studies are heavily biases by inter-subject variation. Recently, we used this framework to create networks matching the morphology of a swine animal model. In appendix EX1 Blood vessel network creation, we create a dataset of this kind and present the analysis.

### 4.2 IFF and drug distribution

To measure IFF and the related interstitial fluid pressure (IFP) in vivo is an especially hard task even with modern measurement methods. Regarding the medical aspects, special interest is drawn to the vasculature as source for therapeutics and nutrients. A solid understanding of IFF and IFP is also crucial for fighting cancer since an elevated IFP is believed to pose a barrier to drug delivery and oxygenation. The resulting hypoxia leads to inefficacy of drugs and radiation. For these type of simulation we consider the IFP as driving force of the flow, i.e. via Darcy’s law. Differences between IFP, blood pressure and the fluid pressure within lymphatics drive transvascular fluxes according to Starling’s equation. We solve the stationary case leading to a Poisson equation for the pressure.

Ensuing the determination of IFF, solute transport into tissue can be simulated. If specified by parameters (see appendix EX3 Interstitial fluid flow and drug transport), the mass balance equation for solutes is integrated numerically based on the calculated flow, diffusion coefficients, rate constants, etc. For an exemplary overview of such quantities, see Fig 5. Quantitatively these agree well with the results obtained by simulating larger systems where the administration of doxorubicin was included. [17, Fig 4,7]. Notably, our results obtained by these simulations suggest that despite of an IFP plateau within the tumor, IFF does not cease and still allows for substantial convective transport which was in contradiction to the view at that time. Furthermore we varied numerous parameters and the administration protocol emulating clinical studies which would be extensive and lengthy in practice.

**Fig 5.**
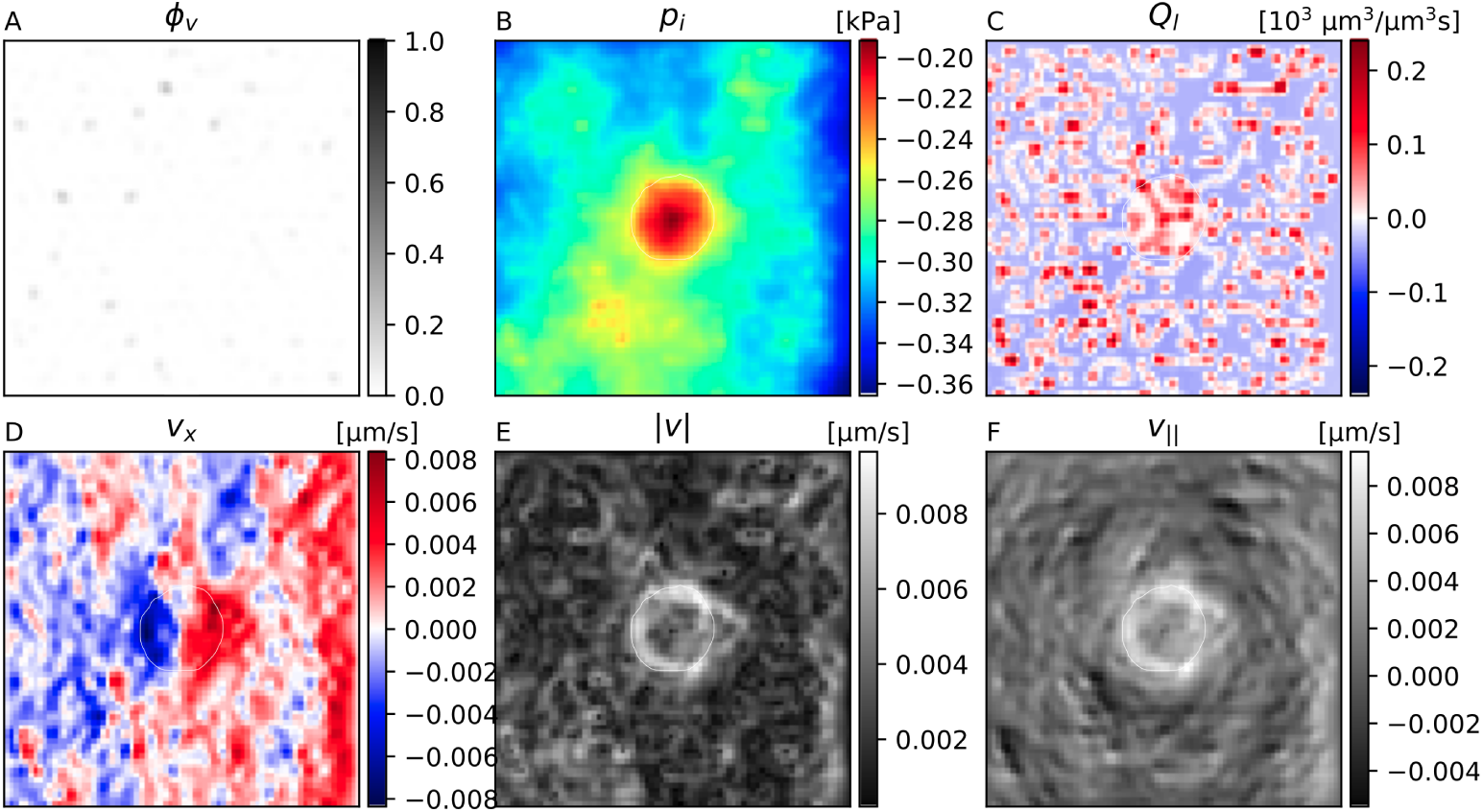
Snapshots of interstitial fluid flow quantities. (A) Vessel volume fraction, (B) IFP, (C) Fluid source term, (D) x-component of the IF velocity, (E) Magnitude of the IF velocity, (F) Projection of IF velocity in outward direction. The plots were generated from 2d slices through the center of the simulation result obtained in appendix EX3 Interstitial fluid flow and drug transport. The contour line indicates the boundary of the viable tumor mass. The internal regions consist of necrotic tissue, while the outer area is normal host tissue.

### 4.3 Detailed oxygen distribution

We augmented the model of oxygen transport such that intravascular variation of the oxygen content along vessel center lines is taken into account. Moreover a varying hematocrit based on the (Red Blood Cell) phase separation effect (see [18] and section Hydro- and hemodynamics,Finite Elements Method) has been implemented. Since most oxygen is transported through the body while bound to the RBCs, the hematocrit distribution is the physical key to the oxygen transport. Together with the unbound oxygen within the blood, we implemented a coarse-grained model of oxygen concentration along the direction of blood flow which causes tissue oxygenation. An example of these type of simulations is presented in supplemental material EX4 Detailed oxygen distribution. Here we show the visualization of the simulation results as Fig 6.

**Fig 6.**
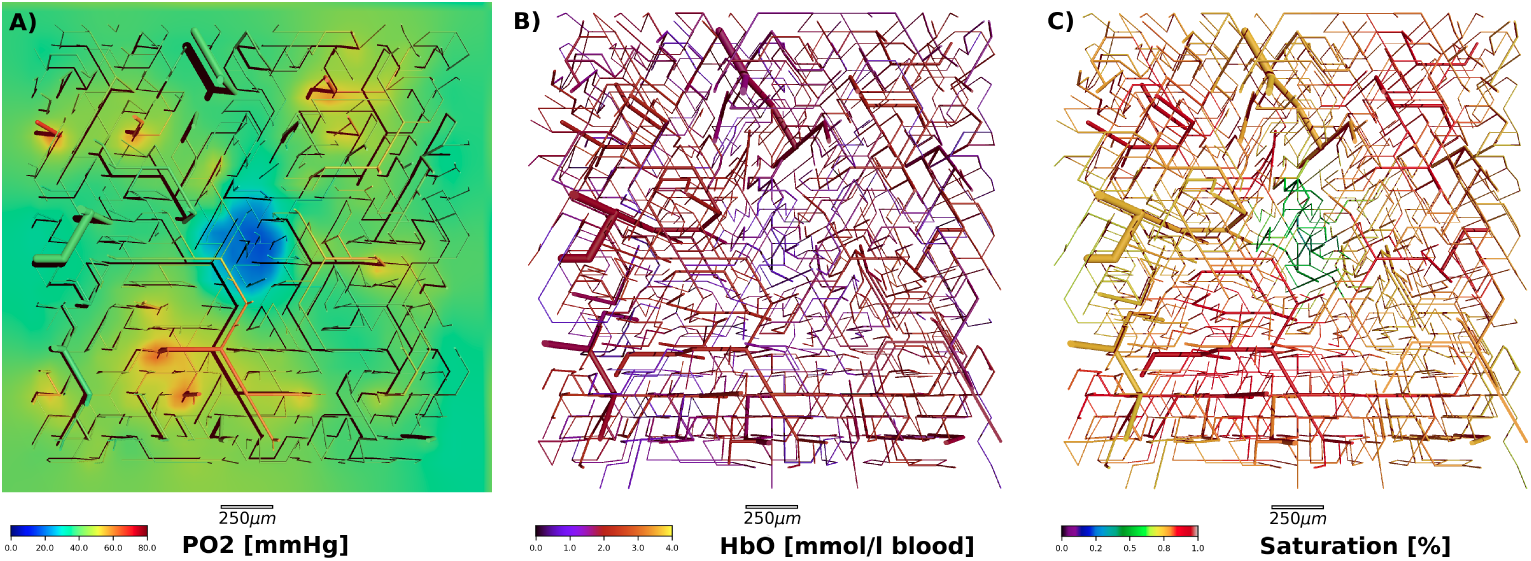
Overview of detailed oxygen simulation. *Inset A)* partial oxygen pressure within vessels and surrounding tissue. Note the hypoxic core in the center where the tumor grows. *Inset B)* amount of oxygenated hemoglobin within the vessels. For illustration purposes the tissue is not depicted. *Inset C)* blood oxygen saturation within the vessels. Note that all three figures show the same blood vessel network. See appendix EX4 Detailed oxygen distribution for construction details.

The focus of the published study was to simulate tissue total hemoglobin concentrations cHb and tissue blood oxygen saturation Y of tumors and host tissue and compare the results with clinical data on breast tumors and surrounding normal breast tissue obtained from a cohort of 87 breast cancer patients using optical mammograph. Following the workflow of *tumorcode* we began by constructing 90 distinct host vasculatures followed by the growth of fake tumors. Using the detailed oxygen simulation, we showed that the redirection of the hematocrit during the tumor growth is accompanied by changes in flow rates and identified this as key element of the modulation in blood oxygen saturation. In contrast, the fact that vascular dilatation leads to an elevated tumor oxygen saturation is not intuitive but confirmed by our simulations. Strikingly we found evidence that different types of initial vascular configurations are correlated to the variance of tissue hemoglobin concentration, tissue blood oxygen saturation and perfusion which raises biological insight.

## 5 Conclusion

This manuscript describes the software used to draw conclusions about elevated interstitial fluid pressure, drug and oxygen distribution in previous publications. All simulations are based on constructing synthetic arterio-venous blood vessel networks on macroscopic tissue volumes matching topological, morphological and hydrodynamic properties observed in real samples. To the best of our knowledge there is no comparable software for this task.

Development, implementation and debugging of software is tedious, and distracts from biological or medical aspects. By varying the parameters in our simulation framework, we allow everybody to tackle its own questions within the realm of this software. Further, we allow everybody to reproduce the experiments we carried out so far, as it is required by good scientific practice. We share the source-code under open source license at http://github.com/thierry3000/tumorcode

Despite the code is platform independent, successful compilation and usage as described in this article is restricted to Unix-based systems at the moment. If required, support for different operating systems is feasible. Likewise, implementing a graphical user interface as extension of our module-based code is straight forward and could serve other researchers in life science with less background in computer science. Following the direction towards a common code base in the field, we hope this article and mainly the examples in the supplemental material help to overcome technical struggles. In future, merging tumorcode with other codes may be an option.

Currently, the drug as well as the detailed oxygen distribution is calculated at time snapshots of the tumor growth process only. To engage pharmacodynamics and improve predictions, correlating the time points could be considered in future. Also additional features relevant for clinics like, pH, ATP consumption or formation of metastasis could be included. The questions in the broad field of cancer research are immense.

## 6 Supplemental

**S1 File Data created in the examples** We add all data produced in EX1 Blood vessel network creation to EX5 Visualization tools.

**S1 Video. Blood vessel network creation process.** To illustrate the process of network creation, we show an example in fast motion. The resulting dataset is quite big and therefore all intermediate stages are omitted. The complete source used to construct the data for the movie can be found in the file *povrayRenderVesselgeneratorDbg.py* for references.

**S2 Video. 3D model of tumor-induced angiogenesis.** To illustrate the tumor model we show a video of the dynamical growth process.

### EX1 Blood vessel network creation

In this supplementary example we create an artificial blood vessel network using *submitVesselgeneration* and show some measurement tools. The algorithmically description is documented in [2].

### EX2 Tumor Growth

In this supplementary example we grow tumors on the previous created artificial blood vessel networks and show how to invoke some quantifications.

### EX3 Interstitial fluid flow and drug transport

Providing a tumor simulation, we describe here how interstitial fluid pressure and drug distributions could be calculated and visualized. These simulation where used for the publication [17].

### EX4 Detailed oxygen distribution

Providing a tumor simulation, we calculate the detailed oxygen distribution as described in [15] and run the scripts in order to analyze those oxygen distribution.

### EX5 Visualization tools

Using the data created in previous examples, we show how the simulation results can be visualized using your tools facilitated by Povray and ParaView.

